# Representations of local spatial information in the human medial temporal lobe during memory-guided navigation

**DOI:** 10.1101/2020.11.18.389346

**Authors:** Shao-Fang Wang, Valerie A. Carr, Serra E. Favila, Jeremy N. Bailenson, Thackery I. Brown, Jiefeng Jiang, Anthony D. Wagner

## Abstract

The hippocampus (HC) and surrounding medial temporal lobe (MTL) cortical regions play a critical role in spatial navigation and episodic memory. However, it remains unclear how the interaction between the HC’s conjunctive coding and mnemonic differentiation contributes to neural representations of spatial environments. Multivariate functional magnetic resonance imaging (fMRI) analyses enable examination of how human HC and MTL cortical regions encode multidimensional spatial information to support memory-guided navigation. We combined high-resolution fMRI with a virtual navigation paradigm in which participants relied on memory of the environment to navigate to goal locations in two different virtual rooms. Within each room, participants were cued to navigate to four learned locations, each associated with one of two reward values. Pattern similarity analysis revealed that when participants successfully arrived at goal locations, activity patterns in HC and parahippocampal cortex (PHC) represented room-goal location conjunctions and activity patterns in HC subfields represented room-reward-location conjunctions. These results add to an emerging literature revealing hippocampal conjunctive representations during goal-directed behavior.

## Main text

The cognitive map hypothesis posits that animals construct structured knowledge of environments to guide behavior (Tolman 1948). This structured knowledge is thought to depend, in part, on two important mechanisms of the hippocampus (HC): (1) conjunctive coding for individual episodes enables distinct features of an event to be bound into a neural representation (e.g., Davachi, Mitchell, & Wagner, 2003; Eichenbaum & Cohen, 2014; Eichenbaum, Dudchenko, Wood, Shapiro, & Tanila, 1999; Schiller et al., 2015); and (2) mnemonic differentiation across multiple episodes mitigates interference between similar events (e.g., Favila, Chanales, & Kuhl, 2016; LaRocque et al., 2013; O’Reilly & Mcclelland, 1994; Yassa & Stark, 2011). The contributions of these two HC computations to spatial navigation have been explored extensively in rodents, but to a much lesser extent in humans.

In rodents, neurons in HC and medial temporal lobe (MTL) cortical regions encode spatial information, as evidenced by place (O’Keefe, Dostrovsky, & J. O’Keefe, 1971), head direction (Taube, Muller, & Ranck, 1990), border (Solstad, Boccara, Emilio, Moser, & Moser, 2008), and grid cells (Hafting, Fyhn, Molden, Moser, & Moser, 2005). Consistent with HC’s fundamental role in conjunctive encoding, HC representations bind non-spatial and spatial features of an environment (Manns & Eichenbaum, 2009; Moita, Rosis, Zhou, LeDoux, & Blair, 2003; Wood, Dudchenko, & Eichenbaum, 1999). For example, neurons in rodent HC subfields CA1 and CA3 encode the conjunction of odors and positions (Komorowski, Manns, & Eichenbaum, 2009). Evidence from pattern similarity analysis further indicates that rodent HC neurons exhibit hierarchical coding — from the overall context to object position, reward, and item identity — that parallels the hierarchical spatial information experienced during the stimulus sampling epoch (McKenzie et al., 2014).

While HC representations are conjunctive within single memory episodes, HC representations differentiate similar environments across different episodes. Within HC, spatial information of different environments are encoded by separable cell assemblies and with distinct firing rates (Colgin, Moser, & Moser, 2008; Lee, Yoganarasimha, Rao, & Knierim, 2004; S. Leutgeb et al., 2005; Vazdarjanova & Guzowski, 2004). More specifically, neuronal ensembles in distinct HC subfields respond differently to changes in the spatial environment, which influence neural representations linearly in CA1 and sigmoidally in CA3. In addition, sparse coding in the dentate gyrus (DG) leads to increased sensitivity to small changes in the environment (J. K. Leutgeb, Leutgeb, Moser, & Moser, 2007; Treves & Rolls, 1994). These findings and a wealth of other data indicate that the mnemonic computations of the rodent HC support the conjunctive coding of spatial information, distinguish different spatial environments or events, and enable present and future memory-guided navigational behavior.

Similar to rodents, place cell-, grid cell-, and head-direction cell-like activity has been found in human HC, entorhinal cortex (ERC), and other brain regions (e.g., Baumann & Mattingley, 2010; Doeller, Barry, & Burgess, 2010; Ekstrom et al., 2003; Epstein, Patai, Julian, & Spiers, 2017; He & Brown, 2019; Jacobs, Kahana, Ekstrom, Mollison, & Fried, 2010; Shine, Valdés-Herrera, Hegarty, & Wolbers, 2016). Findings from early neuroimaging studies indicate that HC activation increases when healthy participants retrieve spatial information (Maguire et al., 1998; Maguire, Frackowiak, & Frith, 1997). Recent univariate functional magnetic resonance imaging (fMRI) studies of human spatial navigation have further revealed that activity levels in HC tracks the topology of a virtual street network (Javadi et al., 2017), and activity levels in both HC and ERC track distances to goals and future behavioral choices (Brown, Hasselmo, & Stern, 2014; Howard et al., 2014). This evidence suggests that human HC and MTL cortical regions also play a role in spatial navigation.

Advances in multivariate pattern analysis (MVPA) enable investigations of information coding in the human brain, providing an opportunity to measure content coding (Haxby et al., 2001; LaRocque et al., 2013; Liang, Wagner, & Preston, 2013; Norman, Polyn, Detre, & Haxby, 2006), memory retrieval (e.g., Gagnon, Waskom, Brown, & Wagner, 2019; Gordon, Rissman, Kiani, & Wagner, 2014; Johnson, McDuff, Rugg, & Norman, 2009; Polyn, Natu, Cohen, & Norman, 2005; Staresina, Fell, Do Lam, Axmacher, & Henson, 2012), and prospection (e.g., Brown et al., 2016; Chadwick, Jolly, Amos, Hassabis, & Spiers, 2015). Studies combining MVPA with virtual navigation tasks indicate that human HC activity patterns reflect conjunctive information of space and time for objects encountered in an environment during navigation (Deuker, Jacob, Schro, & Doeller, 2016). Consistent with HC’s role in differentiating memories for episodic events, other data indicate that HC activity patterns distinguish target locations within a virtual room (Hassabis et al., 2009) and that orthogonalization of HC activity patterns is associated with differentiating ambiguous environments or routes (Chanales, Oza, Favila, & Kuhl, 2017; Kyle, Stokes, Lieberman, Hassan, & Ekstrom, 2015) and buildings/rooms that share overlapping associated task demands (Jiang et al., submitted). Within HC, a recent study revealed differential contributions of human CA3/DG and CA1 in representing episodic contexts (videos) and spatial environments (virtual houses) (Dimsdale-Zucker, Ritchey, Ekstrom, Yonelinas, & Ranganath, 2018). Activity patterns in CA1 for objects encountered in the same video and same house were more similar than for objects from different videos and same/different houses. The opposite effect was found in CA3, such that activity patterns differentiated objects encountered in the same video and same house. Finally, emerging evidence indicates that HC activity patterns code for future goal and sub-goal locations during planning periods prior to navigation (e.g., Brown et al., 2016; Brown, Gagnon, & Wagner, submitted). While these and other studies provide ample evidence that human HC, equipped with its computational abilities to generate multidimensional mnemonic codes, is involved in coding for spatial information and differentiating similar environments, few human studies have explored the interaction between these two HC mechanisms in the context of spatial navigation.

Motivated by extant findings, we examined how neural patterns in the human HC and surrounding MTL cortical regions establish structured knowledge of an environment, including spatial and non-spatial elements, during navigation. We hypothesized that activity patterns in HC will combine both hierarchical spatial structure and reward information given prior evidence of conjunctive coding in HC. Additionally, aligned with studies that have found mnemonic differentiation in HC, we hypothesized that orthogonalized activity patterns in HC would distinguish similar environments. In MTL cortex, we hypothesized that parahippocampal cortex (PHC), relative to perirhinal cortex (PRC) and ERC, would code for spatial features given extant findings in the literature (e.g., Aguirre, Detre, Alsop, & D’Esposito, 1996; R. Epstein & Kanwisher, 1998; Hassabis et al., 2009; Maguire et al., 1997). Finally, because conjunctive coding and mnemonic differentiation are thought to differentially occur within HC, we expected lesser degrees of conjunctive coding and mnemonic differentiation in MTL cortical regions.

To test these hypotheses, here we investigated how global contexts, context-location conjunctions, and rewards are encoded in human HC and surrounding MTL cortical regions. In the study, participants completed a two-day virtual navigation task in which they first learned reward-location associations in two different virtual rooms (Day 1, non-scanned). We then tested their memory for the associations one day later (Day 2) while participants underwent high-resolution fMRI (hr-fMRI). The structure of the environments consisted of global contexts (rooms), local positions (goal locations within each room), and their associated rewards (Figure 1A-B). The two virtual rooms had different shapes (square/circular) and wall colors (blue/green), but shared other features (i.e., area of the rooms, ceiling and floor colors). The same five pieces of furniture were placed along the edges of the two rooms, in the same layout (Figure 1A). Each room contained four hidden goal locations in each of four quadrants, and each location was associated with a letter name (A-D). Across the two rooms, goal locations within a given quadrant differed, as did the letter names assigned to the locations. Within each room, two monetary values ($0.10, $2.00) were assigned across the four locations; there were two low-reward locations and two high-reward locations.

**Figure 1.**
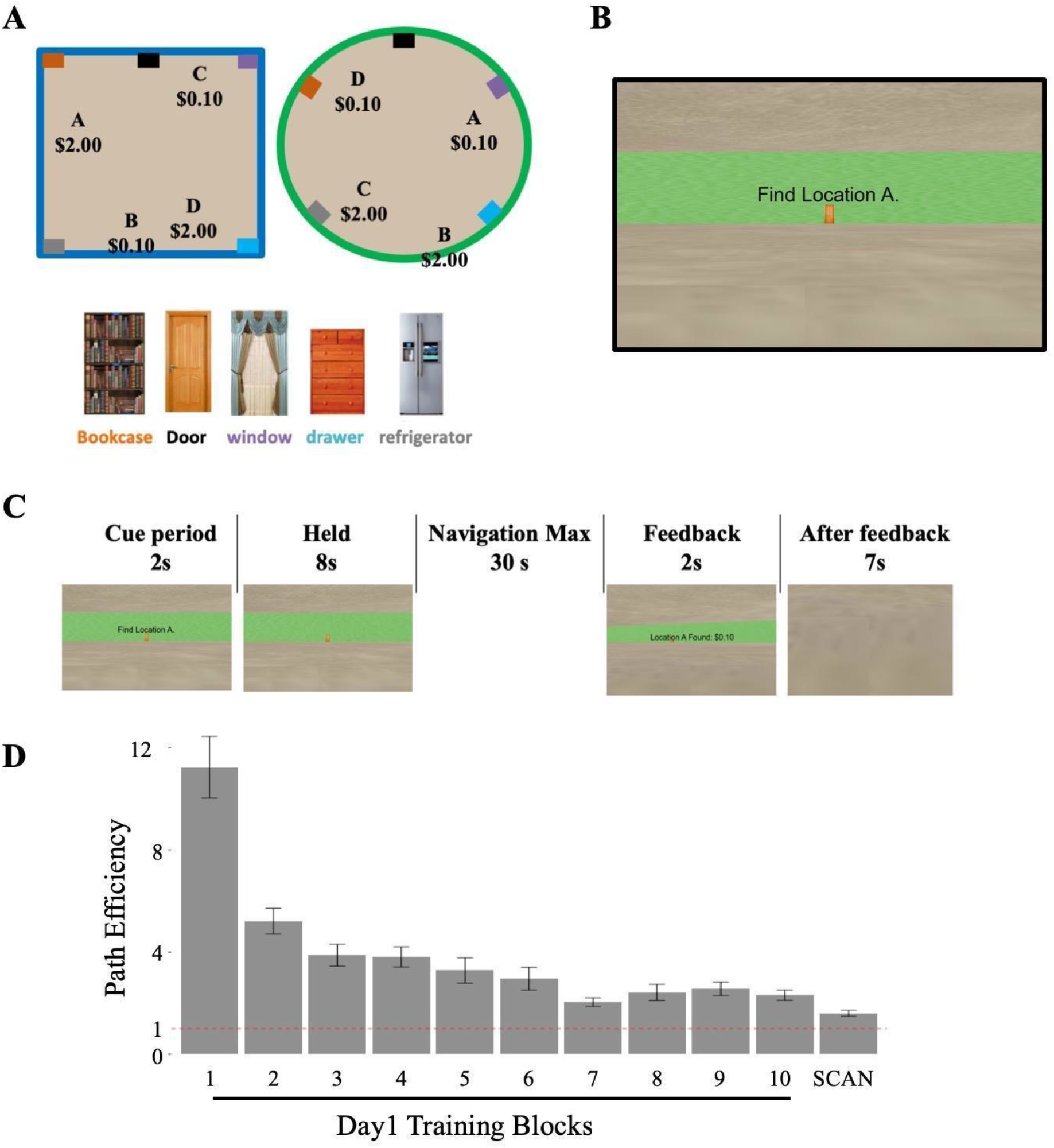
A) Topographic view of two virtual reality rooms and five landmarks in the rooms. One room was square and the other was circular. In this example, the square room had blue walls and the circular room had green walls (color assignment counterbalanced across subjects). Locations A-D within the rooms indicate the hidden goal locations; each was associated with a high or low reward value. Five identical landmarks were placed along the edges of the rooms in the same configuration. Colors of the furniture names below the furniture pictures correspond to the five color boxes in each room in panel A for their locations in the rooms. B) Participants’ first-person view of the room at the beginning of a navigation trial (i.e., cue) on Day 2 inside the scanner. This is an example from the green circular room. Gray ceiling and floor were the same for both rooms. On each trial, participants started in the center of a room facing the wooden door and the goal cue (e.g., “Find location A”) was presented for 2s. C) One representative navigation trial on Day 2 inside the scanner. Periods of interest are cue, held, feedback, and after feedback. There were 12 scan runs, with 12 navigation trials per run (6/room). D) Path efficiency scores for the 10 training blocks on Day 1 and path efficiency scores for the scan session on Day 2. Red dashed line marks path efficiency equals to 1 (i.e., best possible performance). Error bars: across-subject standard error of the mean.

On Day 1, participants (N = 15) completed training tasks designed to foster and validate the learning of the goal locations in each of the two rooms (see Detailed Methods). On Day 2, participants first completed a “top-up” phase to remind them of the rooms, locations, labels, and associated monetary values. After the “top-up” phase, participants completed 12 scan runs during which they were instructed to navigate to cued goal locations in each room while undergoing hr-fMRI. Each fMRI run contained two mini-blocks and each mini-block contained six navigation trials in a room; participants thus completed 72 navigation trials for each room across scanning. During the scans, participants were presented with a text cue for the relevant room (square/circular) before the start of each mini-block. Participants began each navigation trial (Figure 1C) at the center of the room, facing the same piece of furniture — a door — and were presented with a text cue of the name of the goal location for 2 s (cue period). Following the cue period, participants were held at the start location without the goal cue for 8 s (held period) before starting navigation. Participants were then free to navigate, and if they successfully arrived at the cued goal location within a 30-s time limit, they received a text feedback for 2 s (feedback period) displaying the name of the location and the amount of reward. Next, the camera panned down to face the floor (i.e., a grey screen) for 7 s (after-feedback period) before another navigation trial began. If participants did not arrive at the goal location within the 30-s time limit, the trial ended and the next trial began.

We first examined how well participants learned the goal locations on Day 1 (Figure 1D). The initial learning of the two rooms consisted of 10 blocks of exploration-based navigation, in which participants continued to explore/navigate until they had found all four goal locations within time limits (see Detailed Methods). We quantified navigation performance by computing a path efficiency score, which reflects the total distance traveled to a goal location divided by the minimum linear distance to that location from the starting point. A lower path efficiency score would indicate that a participant has learned the goal location and navigated with efficiency from the starting point (1 = perfect efficiency). Analyses revealed that participants’ path efficiency scores decreased as training progressed on Day 1 (first block mean path efficiency = 11.22, SD = 4.67; last training block mean path efficiency = 2.31, SD = 0.77; F(1,14) = 130.68, p < 0.0001), revealing learning of the goal locations.

To determine whether participants retained their knowledge of the environments from Day 1 to Day 2, we compared path efficiency at the end of Day 1 training to Day 2 fMRI navigation runs (Figure 1D). Relative to performance in the final training block on Day 1, path efficiency was even lower during the Day 2 retrieval runs inside the scanner (mean = 1.61, SD = 0.44; F(1,14) = 10.14, p < 0.01), revealing knowledge retention and modest further learning on Day 2. Furthermore, during the fMRI navigation runs, participants successfully navigated to the goal location within the 30-s time limit on 98% (SD = 0.025) of retrieval trials, with an average navigation duration of 7.56 s (SD = 1.5 s). Path efficiency was comparable between high-reward and low-reward goal locations (F(1,14) = 0.97, p > 0.34), and between the square room and circular room (F(1,14) = 0.13, p > 0.72). Collectively, these results show that participants learned the environments, retained their knowledge on the second day, and navigated to goal locations accurately and efficiently during the fMRI navigation runs. Given the small number of failure trials, fMRI analyses were restricted to trials in which participants successfully navigated to the goal location within the 30-s time limit.

We next examined whether and how activity patterns in HC and MTL cortical regions encode the multiple levels of spatial information likely necessary for successful navigation to the goal locations. We used pattern similarity analysis on trial-level parameter estimates to assess whether activity patterns differentiate the two rooms, goal locations within rooms, and reward values. To calculate single-trial parameter estimates for each of the four periods of interest (i.e., cue, held, feedback, and after feedback; Figure 1C), we generated four GLMs, one for each of the four trial periods. For example, to calculate single-trial parameter estimates for the cue period, cue period activity was modeled with multiple single-trial regressions while the other periods (i.e., held, navigation, feedback, and after-feedback periods) were modeled at the condition-level for each run. The trial-level parameter estimates were extracted from anatomically defined MTL regions of interest (ROIs) and submitted to pattern similarity analysis. Specifically, using each participant’s anatomical T2-weighted images and established procedures, we manually traced ROIs for bilateral HC subfields (CA1, CA3/DG, anterior CA1-3/DG, posterior CA1-3/DG, and subiculum), PRC, PHC, and ERC (Figure S1) (Olsen et al., 2009). From these ROIs, we also created a HC ROI by combining all HC subfield masks. We indexed pattern similarity by correlating the vectors of parameter estimates from trial pairs and generated similarity scores by averaging correlations across all trials for each condition of interest.

Results of the pattern similarity analyses indicate that, among all MTL ROIs (HC, PHC, PRC, and ERC), activity patterns in HC and PHC differentiated goal locations within a room upon goal arrival (Figure 2A & C; for findings in PRC and ERC see Figure S2). Specifically, during the feedback period, pattern similarity scores for navigation trials to same goal locations within the same room were significantly higher than that for different goal locations within the same room in HC, HC subfields, and PHC (MTL ROIs × conditions (same/different locations): condition: F(1,14) = 6.82, p < 0.03; ROIs × conditions: F(3,42) = 3.57, p < 0.03; post-hoc paired t-test: HC: t(14) = 3.10, p < 0.035; PHC: t(14) = 4.2, p < 0.004; Bonferroni corrected. HC subfields ROIs × conditions (same/different locations): condition: F(1,14) = 6.91, p < 0.02; ROIs × conditions: F(4, 56) = 0.44, p > 0.78). A control analysis revealed that the location effect was not driven by letter names of the locations (A-D): pattern similarity scores for across room same and different goal location names did not significantly differ (MTL ROIs × conditions (same/different letter names): condition: F(1,14) = 0.40, p > 0.5, ROIs × conditions: F(3,42) = 0.74, p > 0.5. HC subfields ROIs × conditions (same/different letter names): condition: F(1,14) = 0.39, p > 0.54; ROIs × conditions: F(4, 56) = 0.86, p > 0.49).

**Figure 2.**
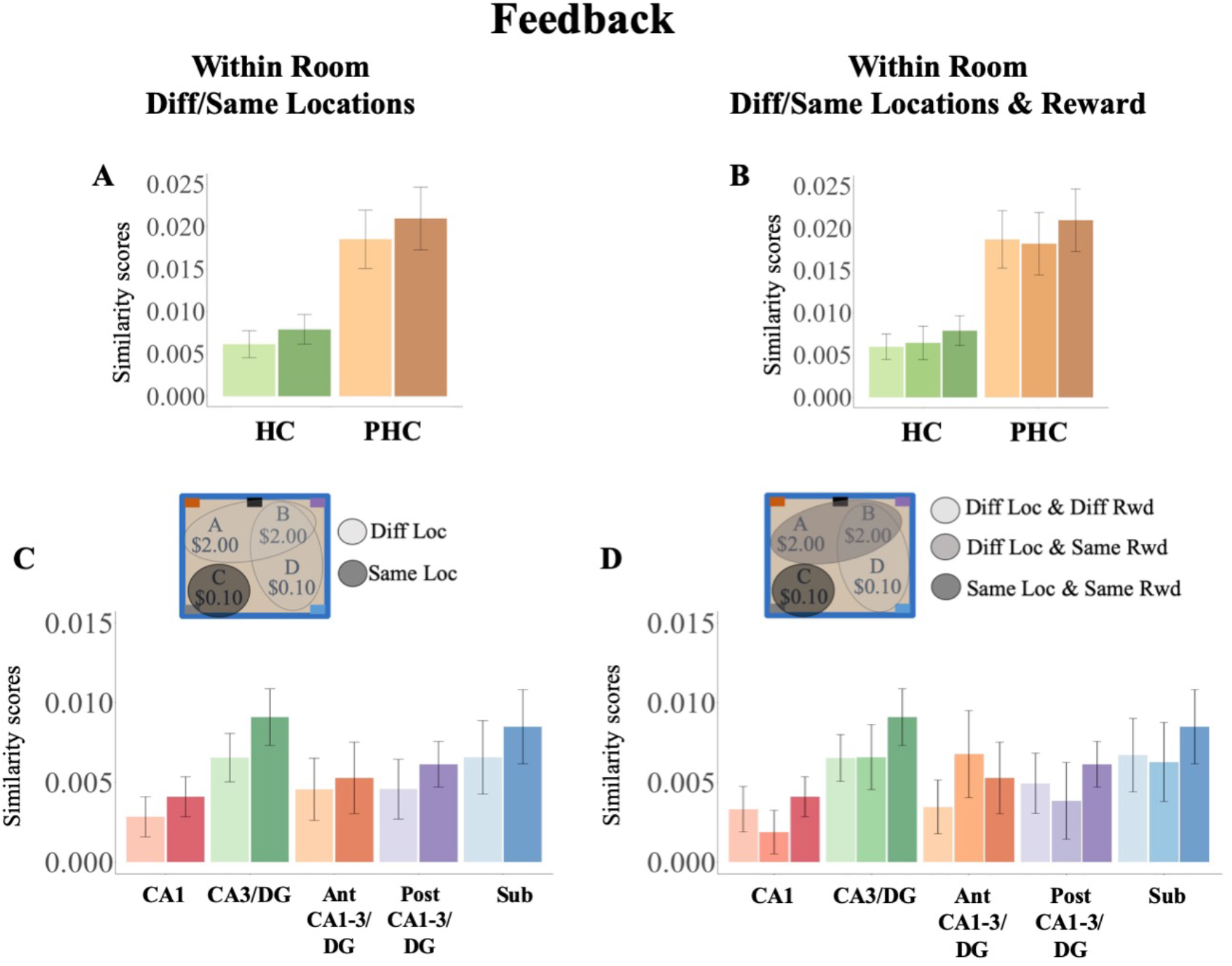
Pattern similarity scores in MTL cortical (A/B) and HC subfields ROIs (C/D) for within-room different/same locations and within-room different/same location and reward during the feedback period. For the plots of within-room different/same locations (A/C), the light color bars represent similarity scores for different locations and the dark color bars represents similarity scores for the same locations. For the plots of within-room different/same locations and reward (B/D), the light color bars represent similarity scores for different locations, the intermediate color bars represent similarity scores for different locations and same reward values, and the dark color bars represent similarity scores for the same location and same reward. Conditions of the bars are demonstrated in the legend: the legends use topographic view of the square room with blue walls as an example similar to Figure 1A. The gray circles in the topographic view of the room are example conditions. Error bars: across-subject standard error of the mean.

We next investigated whether the location effect was influenced by reward-location associations, first examining neural patterns for reward values regardless of goal locations. Results revealed that, within the same room, pattern similarity scores for navigation trials with the same reward value (regardless of whether or not location differed) were not significantly different from those for different reward values (location differed) in both MTL ROIs and HC subfields (MTL ROIs × conditions (same/different reward values): condition: F(1, 14) = 0.06, p > 0.81; ROIs × conditions: F(3,42) = 2.65, p > 0.06; post-hoc paired t-test: ps > 0.1; Bonferroni corrected. HC subfields ROIs × conditions (same/different reward values): condition: F(1,14) = 1.39, p > 0.25; ROIs × conditions: F(4,56) = 1.50, p > 0.21). When combined with the preceding analyses, these results indicate that there were location-specific activity patterns, but not location-free reward patterns, in HC and PHC when participants arrived at the goal locations. Next, we examined neural patterns for different/same reward values and different/same goal locations (Figure 2B & Similarity scores for different locations and different reward values, different locations and same reward values, and same location and same reward values were not significantly different within the same room in both MTL and HC subfields ROIs (MTL ROIs × conditions (same/different locations and same/different reward values): condition: F(2, 28) = 2.68, p > 0.086; ROIs × conditions: F(6,84) = 1.72, p > 0.13; HC subfields ROIs × conditions (same/different locations and same/different reward values): condition: F(2, 28) = 2.28, p > 0.12; ROIs × conditions: F(8,112) = 1.43, p > 0.19). Post-hoc mixed-effect models revealed that, in PHC, similarity scores of same location and same reward values were significantly higher than that for different locations and different reward values (t(28) = 2.69, p < 0.05; Bonferroni corrected) (Figure 2B).

We next performed the above location and reward analyses for the after-feedback period — when participants viewed a grey screen following navigation — and found that activity patterns in the MTL ROIs did not significantly differentiate different locations within the same room, though there was a trend (MTL ROIs × conditions (same/different locations): condition: F(1,14) = 3.81, p > 0.07; ROIs × conditions: F(3,42) = 1.89, p > 0.15; post-hoc paired t-test: ps > 0.07; Bonferroni corrected) (Figure 3A; for findings in PRC and ERC see Figure S2). However, analyses focused on HC subfields revealed a main effect of condition, where pattern similarity scores for the same location were significantly higher than those for different locations within the same room (HC subfields ROIs × conditions (same/different locations): condition: F(1,14) = 6.73, p < 0.03; ROIs × conditions: F(4,56) = 1.30, p > 0.28) (Figure 3C). Again, a control analysis showed that the location effect was not driven by letter names of the locations (A-D) (HC subfields ROIs × conditions (same/different letter names): conditions: F(1,14) = 0.001, p > 0.97, ROIs × conditions: F(4,56) = 2.38, p > 0.06; post-hoc paired t-test: ps > 0.20; Bonferroni corrected). Moreover, there was a significant interaction between HC subfields and pattern similarity scores for same and different reward values within the same room (HC subfields ROI × conditions (within room same/different reward values): condition: F(1,14) = 2.36, p > 0.1; ROIs × conditions: F(4, 56) = 2.78, p < 0.04; post-hoc paired t-test: ps > 0.1; Bonferroni corrected). Finally, there was a significant main effect for same/different locations and same/different reward values (HC subfields ROI × conditions (same/different locations and same/different reward values): condition: F(2, 28) = 4.12, p < 0.03; ROIs × conditions: F(8,112) = 1.33, p > 0.23). Post-hoc mixed-effect models revealed that similarity scores of same location and same reward values were significantly higher than that for the other two conditions (different locations and same reward values; different locations and different reward values) (post-hoc mixed effect model: same location vs. different location and same reward values: t(204) = −2.9, p < 0.004; same location vs. different locations and different reward values: t(204)= −2.7, p <0.008) (Figure 3D). While the findings in the feedback period were qualitatively similar but did not reach statistical significance, these finding indicate that there were conjunctive representations of reward and location in HC subfields during the after-feedback period.

**Figure 3.**
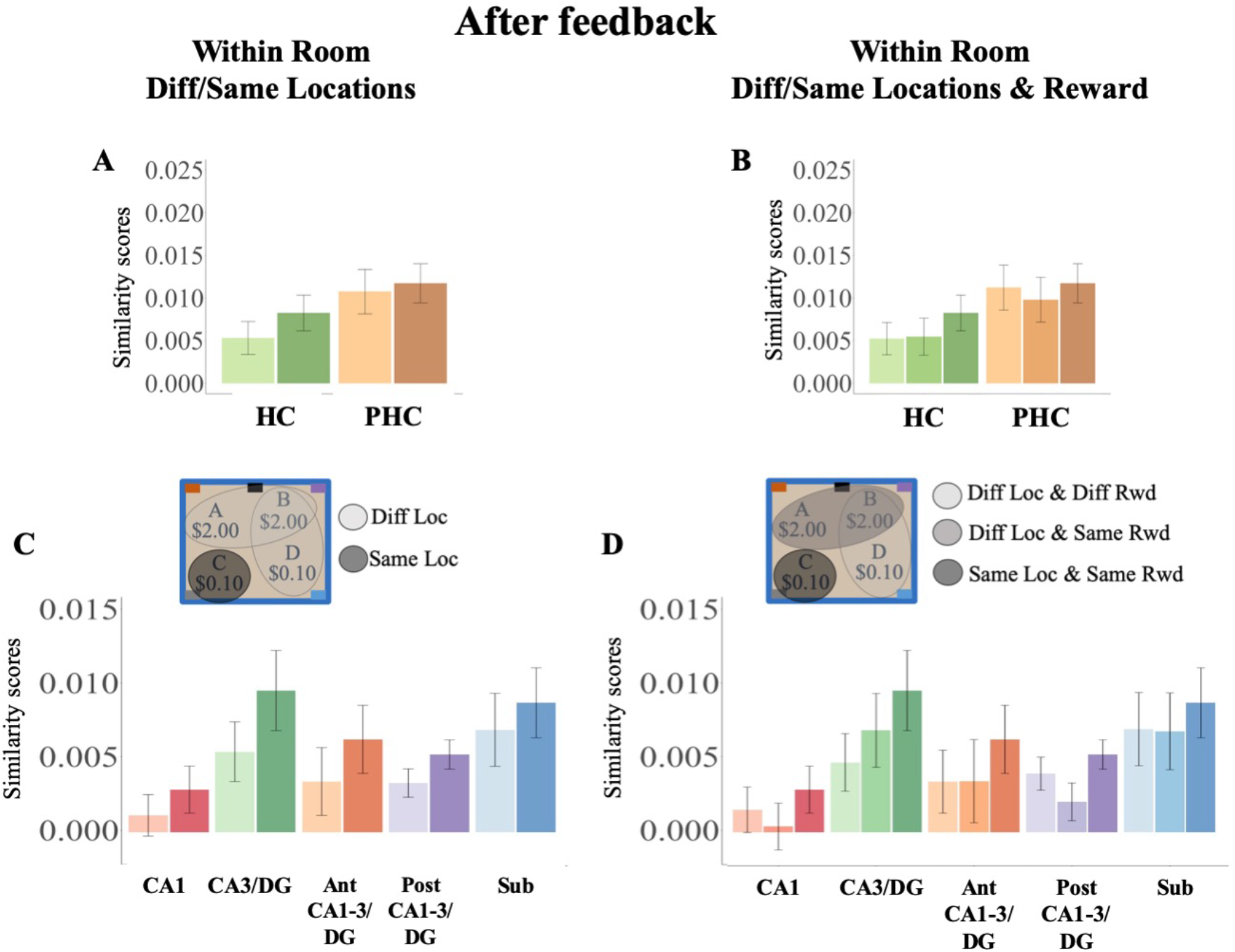
Pattern similarity scores in MTL cortical (A/B) and HC subfields ROIs (C/D) for within-room different/same locations and within-room different/same location and reward during the after-feedback period. Conditions of the bars are the same as Figure 2 and are demonstrated in the legend. Error bars: across-subject standard error of the mean.

Next, we examined whether HC and MTL cortical regions encode other types of spatial features, such as global contexts and reward values, and additionally, whether the neural patterns coded for spatial features within the same room described above were room specific. We computed similarity scores for same and different rooms (i.e., global contexts), across room same/different reward values (i.e., reward values), and across room same/different goal quadrants. The analysis of across room same/different goal quadrants tested whether the within room goal-location effects found during feedback and after-feedback periods were room specific. Pattern similarity scores for navigation trials to goal locations in the same quadrants across the two rooms were “same goal quadrants” and goal locations in different quadrants across the two rooms were “different goal quadrants”. These analyses revealed no significant differences in similarity scores for all of the above conditions during any period (see the Supplement for details). These results suggest that, in contrast to the effects observed within a context/room, activity patterns did not differentiate global contexts (i.e., rooms) and reward values. Thus, the location specific activity patterns (a) in HC and PHC during feedback and (b) in HC subfields during after-feedback periods were room specific and (c) the reward-location activity patterns in HC subfields during after-feedback period were also room specific.

To this point, the results reveal room-location neural patterns in HC and PHC during feedback and room-reward-location neural patterns in HC subfields after feedback. The temporal differences between these two effects may be due to differential sensory and psychological inputs during these two stages of a trial. During the feedback period, participants were at the goal location with a view of the room from the goal location and text feedback. During the after-feedback period, the camera panned down to face the floor (i.e., a grey screen). We designed the after-feedback period in this way to control for visual inputs across the different goal locations and rooms. Without visual input of the room, it is possible that spatial information was less dominant during after feedback, and therefore, neural activity patterns represented not only spatial information but also its conjunction with reward information. It is also possible that participants were processing and reflecting on the room-reward-location associations during the after-feedback period. Although differences in visual inputs may contribute to the differences in neural activity patterns, we do not think that the room-location neural patterns in HC and PHC during the feedback period were entirely driven by goal location specific views and text feedback. This is because the goal locations fell a minimum distance from the walls (mean distance from the walls 16.57 a.u., SD = 2.67; square room dimension: 150 × 150 a.u.; circular room radius: 84.63 a.u.). Therefore, the view of the room from the goal locations was still dominated by features shared across goal locations, such as the wall, ceiling, and floor of the room. In addition, the control analysis revealed that the location effect was not driven by letter names of the locations, excluding the possibility that feedback text was driving the location effect. However, further studies are required to test whether neural activity patterns in HC are modulated by such sensory inputs.

The lack of differential activity patterns for the global contexts (i.e., the two rooms) may be due to an overtraining effect for our participants. With a 98% (SD = 0.025) success rate and highly efficient navigation during scanning (mean path efficiency = 1.61, SD = 0.44), participants were very familiar with the eight goal locations and may have treated each location independently. This observation is supported by evidence from rodent spatial learning studies. In one study, as learning progressed, the number of rodent hippocampal neurons that encoded order-position associations increased, and, more importantly, the encoding selectivity of the neurons also increased from responding to all positions in a context to position-specific responding in the context (Komorowski et al., 2009). It is possible that, had we collected fMRI data while participants learned the two virtual rooms on Day 1, neural activity patterns would have reflected the global contexts. As learning progressed, the global contexts may have become less critical for navigating as participants became more familiar with the differences across the two rooms. Additional studies are needed to characterize the ways in which neural activity patterns in human HC and MTL cortical regions change across learning and are updated by experience.

Findings from the current study are largely consistent with results from prior human spatial memory studies in showing that activity in CA3/DG encodes conjunctive spatial information and differentiates different environments either during the learning of the environments or the retrieval of specific details of learned environments (e.g., Brown et al., 2014; Dimsdale-Zucker et al., 2018; Kyle et al., 2015; Stokes, Kyle, & Ekstrom, 2015). Unlike prior studies, however, our results did not show differences among HC subfield representations. The recent studies from Stokes et al. (2015) and Kyle et al. (2015) demonstrated that CA3/DG activity patterns were more sensitive to differences in spatial environments than CA1 patterns. These divergent results in subfield functions may be due to important design differences between the current study and prior studies: 1) In this study, there were no ambiguous environments created to induce interference, and 2) the participants in this study learned the environments very well before undergoing retrieval inside the scanner. For both prior studies, researchers created ambiguous spatial environments using the geometry or details within each environment which may have induced a high demand for the memory system to differentiate these environments. In our design, the two virtual rooms had different shapes, wall colors, and explicit labels so there was less of a demand for discriminating between the two environments. Furthermore, the fact that the current study and previous studies focus on representations gathered from different stages of the learning process may also contribute to the differences in findings.

The results presented here are also largely consistent with a number of predictions from theoretical models of episodic memory. It is widely believed that neurophysiological characteristics of HC subfields give rise to HC’s abilities to distinguish similar episodes during encoding and to retrieve previously stored episodes using partial cues (Amaral, 1993; O’Reilly & McClelland, 1994; Treves & Rolls, 1994; Yassa & Stark, 2011). More specifically, computational models and anatomical tracing studies suggest that sparse neural activity in DG and its connection with CA3 allows them to re-code overlapping cortical inputs for an episode in a compressed format. This orthogonalized activity pattern of the episode reduces interference between traces. Furthermore, recurrent connections in CA3 enable reinstatement of previously learned activity patterns when a partial cue is presented. Reinstated information in CA3 is transmitted back to the cortex via CA1 and subiculum, so the representations in CA1 could be strongly influenced by those in CA3, especially during reinstatement. This is consistent with our results showing no differences between CA3/DG and CA1 conjunctive representations during retrieval. Furthermore, some models based on rodent spatial studies suggest that both CA3 and CA1 can perform pattern separation when inputs are dissimilar (Guzowski, Knierim, & Moser, 2004). This may be the case in our study since (1) our environments are perceptually very different and (2) participants are over-trained and can distinguish the two environments very well.

Taken together, our results demonstrate that HC and PHC encode specific room-location conjunctions and HC subfields encode room-reward-location associations during navigation within familiar environments. These findings add to the literature documenting that HC forms conjunctive and differentiated representations of local spatial information during memory-guided navigation.

## Acknowledgments

This project was supported by grants from the National Institutes of Health (R01-MH076932; R21-AG058111), the Marcus and Amalia Wallenberg Foundation (Marcus Wallenbergs Stiftelse för Internationellt Vetenskapligt Samarbete), and a Government Scholarship to Study Abroad from the Ministry of Education, Taiwan. We thank Shaw Hsu and Corey Fernandez for helpful comments and discussions on findings and drafts of the manuscript, and Madison Hunt for helpful input.

The data that support the findings of this study are available from the corresponding author upon reasonable request.

## Detailed Methods

### Participants

Twenty-seven right-handed, healthy adult participants were recruited from the Stanford University community and surrounding area. Data from 12 participants were excluded from analyses (motion sickness: 1; excessive head motion: 6; MR spiking artifacts: 5). For the 15 participants included, mean age was 22.5yrs (range 19-25 yrs; 3 females). All participants had normal or corrected-to-normal color vision. Informed written consent was obtained from all participants, who were paid for their participation in a manner approved by the Stanford University Institutional Review Board.

### Behavioral paradigm

#### Virtual environments

The behavioral task required participants to navigate in two different virtual environments/rooms, using cues to navigate to four learned hidden locations in each; each location was associated with one of two reward values (Figure 1). The two virtual rooms were created using Vizard 3.0 (WorldViz; http://www.worldviz.com/products/vizard). One room had a square base and the other had a circular base; the area of the rooms was identical (square room dimension was 150 × 150 arbitrary unit (a.u.); circular room radius was 84.63 a.u.). Room walls were either green or blue; wall color was counterbalanced across subjects. The floor and ceiling of both rooms were grey. To facilitate learning and memory-guided navigation, the same five distinct landmarks were placed along the edges/walls of both rooms in the same configuration; landmarks were sized at 2 × 2 a.u.. Four of the landmarks — a refrigerator, a bookcase, a full-length window with curtains, and a chest of drawers — appeared in the four corners of the square room, and at the corresponding locations in the circular room. The fifth landmark — a door — appeared at the center point along one wall of the square room, and at the corresponding location in the circular room. We used the same landmarks and the same configuration of the landmarks across the two room to rule out the possibility that neural activity pattern reflects specific furniture near each goal location, not the actual spatial location.

To place the centers of the goal locations, we generated four sets of four goal locations (one for each quadrant) (Figure S3). Goal locations in each set were positioned such that (a) there was one location per quadrant in each room; (b) the distances between goal locations to the room center were around 40 a.u., 50 a.u., 60 a.u. and 70 a.u. (mean +/− SD: 40.25 +/−1.94 a.u.; 51.53 +/− 0.51 a.u.; 60.06 +/− 0.83 a.u.; 69.3 +/− 1.6 a.u.); (c) the locations fell a minimum distance from the walls (mean distance from the walls 16.57 a.u., SD = 2.67), and a minimum distance to vertical midline (y = 0) (18.75 a.u.) and horizontal midline (x = 0) (19.5 a.u.). The mean distance between goal locations within a room was 89.16 a.u. (SD = 22.3 a.u). For each subject, two out of the four sets of goal locations were used as goal locations in the virtual rooms (one set for each room) such that the locations were non-overlapping across the two rooms. Participants’ arrival within a 5 a.u. radius from the center of the goal location was considered a successful trial. Two locations were associated with a low monetary value ($0.10) and two with a high value ($2.00). To enable cueing of navigational goals on a trial-by-trial basis, each location was associated with an alphabetic name (‘A’,’B’,’C’,’D’). The reward-name-location labeling varied across the two rooms, with the constraints that at least one reward-name-location combination (e.g., $2.00 for location A in quadrant one) was different across the two rooms (10 subjects had all eight different combinations; 5 subjects had six different combinations: one reward-name-location combination was the same across the two room). The sets of goal locations used, their assignments to the square/circular room, and the color of the rooms were varied across subjects.

We created a practice room to permit navigational training at the outset of the experiment. The practice room was rectangular in shape with brown walls, had no landmarks, and was substantially smaller than the square and circular rooms (50 ×25 a.u.). One hidden location was placed in the second quadrant of the practice room, with a distance to room center of 20.62 a.u. and a distance to the nearest wall of 4.5 a.u..

During training outside of the scanner, the arrow keys on a standard keyboard were used for virtual navigation. The up arrow and down arrow keys supported forward and backward movement; the left and right arrow keys supported counterclockwise and clockwise rotation. Inside the scanner, four buttons on a response box corresponding to the arrow keys were used for virtual navigation.

### Task design

Day 1: training and testing (~2.5 hrs)

During Day 1, participants performed a series of computer tasks to allow them to learn the locations of the hidden goals and their associated labels and rewards in the two rooms. Day 1 began with a short practice phase, which familiarized participants with the keys to navigate. During the practice phase, participants navigated in a small rectangular virtual room to look for one hidden reward over four trials. The practice room was purposefully different from the experimental rooms to avoid familiarity effects. For each practice trial, participants had 3 min to find the goal location; if they found the goal location before the time was up, they were permitted to navigate freely in the room. When arriving at the goal location, text feedback was given (“You found the sample reward. Good job!”). This feedback differed from that used in the experimental task.

Subsequently, to enable initial learning of the two rooms and their goal locations, participants completed a 10-run exploration-based learning phase followed by a 4-run cue-based testing phase for each room. For the 10-run exploration-based learning phase, each run consisted of four trials in the same room. Each run began with the participant in the middle of a room; participants were instructed to freely navigate in the room to find the four hidden goals. During this phase, when the participant encountered each hidden location while exploring, a label for the location (A, B, C, or D) and the location’s value was presented for 3 s. During this exploration-based learning phase, participants were to continue to explore/navigate until they had found all four goal locations within time limits (i.e., participants were not returned to the center of the room after finding a location, but instead navigated from one found location to the next location to be discovered/remembered). Critically, the time limits decreased as run number increased (3 min for run 1 and 2; 2 min for run 3 and 4; 1.5 min for run 5 and 6; 1 min for runs 7 to 10). After each 10-run learning phase for a room, participants’ knowledge of the locations was tested (four runs; six trials/run) using a cueing procedure. Specifically, on each trial, participants began the trial at the center of the room facing the door and were cued to navigate to a specific location (e.g., “Find Location A”). Cues were presented for 2 s, and then participants were held at the start location without the cue for 2 s before they could initiate navigation. Feedback was provided when the participant successfully arrived at the cued location for 2 s (e.g., “Location ×found:$X”). Each participant first learned and was tested on the locations for one of the rooms (mean = 39.71 min; SD = 11.11 min), and then learned and was tested on the second room (mean = 33.5 min; SD = 7.38 min). The order of room learning (square/circular vs. circular/square) was counterbalanced across participants.

Following these learning and test phases, there was a final interleaved-testing phase (four runs; six trials/run) (mean = 17.07 min; SD = 2.95 min) designed to be similar to the scanned version of the task on Day 2. For a given run, participants performed six cued navigations in one room, with each trial starting from the room center; runs alternated between rooms. In total, participants performed 12 navigation trials (two runs) in each room, which entailed cued navigation to each of the four goal locations three times. At the beginning of each run, participants received a 4-s room cue (“Moving to the square room” or “Moving to the circular room”), followed by a 3-s delay (fixation cross on the screen) before the first trial of the block started. If a participant did not find the location within 60 s, feedback (“You have run out of time to find this location”) was presented for 2 s and the next trial started. If a participant found the cued location, feedback (“Location ×found: $X”) was presented for 2 s. There was a 4 s inter-trial interval, during which a fixation cross was presented.

Reward values associated with goal locations translated to bonus payment depending on successful navigation to goals. We randomly selected one third of the trials from the interleaved-testing phase as bonus payment in addition to base payment ($10/hr). If participants successfully arrived at the goal locations for the selected trials, we paid them according to the reward associated with the goal locations. The maximum amount of bonus for each participant was capped at $8 (mean = 7.74; SD = 0.54).

Day 2: testing and fMRI scanning (~3.5 hrs)

On Day 2, hr-fMRI data were collected while participants navigated to cued goal locations in each room. Prior to scanning, there was a “top-up” phase designed to remind participants of the rooms, locations, labels, and associated monetary values. First, participants freely navigated for four runs in each room to look for the goal locations. Next, participants performed six cued navigation trials in each room, with runs alternating between rooms. This phase continued until the participant reached a performance criterion of being able to find all locations in a given room within 30 s. Once the criterion was reached (mean number of runs = 5.07; range 4-9), the participant advanced to the scanning phase.

During each of 12 fMRI scan runs, participants were cued to navigate to goal locations within each room. Each run consisted of two mini-blocks of six trials, with one mini-block per room, giving a total of 12 trials/run. The design of each mini-block was similar to the final interleaved-testing phase on Day 1. Prior to each mini-block, participants received a 4-s room cue (“Moving to the square room” or “Moving to the circular room”), followed by an 8-s delay with a fixation cross on the screen before the first trial of the mini-block started. Each mini-block had six trials, and included navigating to all four locations at least once, as well as repeats of two locations – a procedure designed to prevent participants from anticipating remaining goals. Construction of trials met the constraint that (a) each location was cued either once or twice within a run, (b) no location was cued in consecutive trials, and (c) each location was cued three times across two runs. On each trial, participants started in the center of a room facing the wooden door, and were cued to navigate to one of the locations. The goal cue (e.g., “Find location A”) was presented for 2 s, which was followed by an 8-s delay during which participants were held at the start location. Subsequently, participants had a maximum of 30 s to navigate to the goal location. If participants found the location, feedback (“Location ×found: $X”) was presented for 2 s. Next, the camera panned down and remained facing the floor for 7 s. If the location was not found within the time limit, feedback (“You have run out of time to find this location”) was presented for 2 s, followed by a fixation cross (4 s) before the next trial started. Unsuccessful trials were not included in analyses. Across the 12 runs, there were 144 trials, with 72 trials for each room and 18 trials for each location within each room.

Similar to the Day 1 interleaved-testing phase, we randomly selected one third of the trials from the 12 fMRI runs as bonus payment in addition to base rate ($20/hr). The maximum amount of bonus for each participant was also capped at $8 (mean = 7.44; SD = 1.24).

### MRI data acquisition

MRI data were acquired on a 3T GE Discovery MR750 system (GE Healthcare) using a 32-channel radiofrequency receive-only head coil (Nova Medical). High-resolution structural images were acquired using a T2-weighted spin-echo sequence (repetition time = 4200 ms; echo time = 66.1 ms; 0.43 × 0.43 × 2 mm; FOV = 220 mm) consisting of 29 contiguous slices, oriented perpendicular to the main axis of the HC to allow for manual segmentation of the HC subfields (CA1, CA3/DG, and subiculum) and MTL cortical regions (PRC, PHC, and ERC). High-resolution in-plane-accelerated EPI functional images (in-plane acceleration factor = 2; repetition time = 2000 ms; echo time = 32 ms; flip angle = 77°; FOV = 220 mm; 1.67 × 1.67 × 1.5 mm) consisted of 27 contiguous slices co-planar with the structural images.

### fMRI preprocessing

In-house scripts in MATLAB (The MathWorks, Inc., USA) and SPM12 (https://www.fil.ion.ucl.ac.uk/spm/) were used for image preprocessing. First, we observed abnormal MR spike artifacts in raw data according to QA reports generated by the MRI facility. To detect the abnormal MR spike artifacts, we calculated z-scored mean signal from each slice of the brain at each TR. We excluded a TR if it contained at least one slice with z-scored mean signal larger or smaller than 6 standard deviations of the slice for the run. We excluded the whole run when the run contained more than 5% of TRs had abnormal MR spike artifacts. Three runs for two participants in total were excluded based on this procedure. Three participants were excluded at this stage because every run of their data had more than 5% of TRs with abnormal MR spike artifacts. The MR-spike-artifacts-removed data were passed to the following preprocessing steps. The Artifact Detection Tools (ART) was used to detect other artifact TRs: TRs with mean global signal greater than 3 standard deviations of the run mean or TRs with framewise displacement greater than 0.5 mm. All TRs with spike artifacts were marked as “contaminated” TRs and excluded from the analysis. Overall, runs with more than 5% of contaminated TRs (spike artifacts, global signal, and framewise displacement) were also excluded. After artifact corrections, participants who had less than six runs of data were excluded from the analyses, resulting in 15 participants contributing data to the final analyses (for these participants: mean = 27.5 of TRs censored = 1.3% of TRs; mean = 1.8 of runs excluded, range of 9-12 runs/participant, with 12 participants having ≥ 11 runs). The first 5 volumes of each scan run were excluded to permit T1 equilibration. Functional data from remaining participants and runs were slice-time and motion corrected.

### Anatomically defined regions of interest

Using established procedures (Olsen et al., 2009), participant-specific anatomical regions of interest (ROI) were manually defined using the structural images (Figure S1). ROIs included left and right HC subfields (CA3/DG, CA1, anterior CA1-3/DG, posterior CA1-3/DG, and subiculum), PRC, PHC, and ERC.

### Pattern similarity analysis

To maintain high spatial resolution, functional data were neither smoothed nor normalized. To estimate voxel-level BOLD activity related to task events, preprocessed imaging data were analyzed using a general linear model (GLM) approach in SPM12 that generated single-trial parameter estimates (2-s epoch, convolved with the canonical hemodynamic response function). Four GLMs were defined, yielding single-trial parameter estimates for each of the four task-periods of interest (i.e., cue, held, feedback, and after feedback). For each period of interest (i.e., one GLM), single-trial regressors were assigned to trials in which participants successfully navigated to goal locations within a 30-s time limit. The rest of the periods were modeled at the condition-level. If a participant successfully navigated to all goal locations and no runs were excluded due to artifacts (see above), there were in total 144 regressors of interest (144 trials in total; mean = 132 trials, SD = 13.13). For example, to calculate single-trial parameter estimates in response to the goal cues, each cue period of a successful trial was modeled by a single regression (2-s epoch), and the held, navigation, feedback, and after-feedback periods were modeled at the condition-level for each run. Temporal delay between each single-trial parameter estimates is 24.56 on average (8 s + 7.56 s + 2 s + 7 s: held + average navigation duration + feedback + after feedback).

For each participant, the trial-level parameter estimates from each ROI were extracted and submitted to the critical pattern similar analyses, which allowed us to examine whether each ROI carried information about room (circular/square), location within a room (A/B/C/D), and value (high/low). Pearson correlations were calculated for all possible pairs of locations across both rooms using trial-level parameter estimates. Critically, pairs were always drawn across runs (i.e., no within-run correlations were included when calculating pattern similarity scores).

Pattern similarity scores were calculated by averaging the correlations for different conditions to understand neural representations of global contexts (rooms), local position (goal locations), and reward values. To examine neural representation of global contexts — regardless of location and reward values — pattern similarity scores for trials within the same room and trials within different rooms were computed. To examine neural representations of local position within and across the global contexts — regardless of reward values — we computed pattern similarity scores for goal locations for same vs. different goal locations within the room (i.e., within room same/different locations), as well as goal locations that were in the same quadrant across rooms (i.e., across rooms same/different quadrants). To examine neural representations of reward values within and across the global contexts — regardless of location (i.e., quadrants) — we calculated similarity scores for within/across room same and different reward values. To examine conjunctive neural representation of spatial location and reward values in the same room, pattern similarity scores were calculated for within room same location and same reward values, within room different locations and same reward values, and within room different locations and different reward values (within room different/same locations & reward). Finally, we also performed a control analysis to see whether neural representations encoded the goal locations with same and different letter names (A-D) across rooms.

## SUPPLEMENTAL MATERIALS

### Supplemental figures

**Figure S1.**
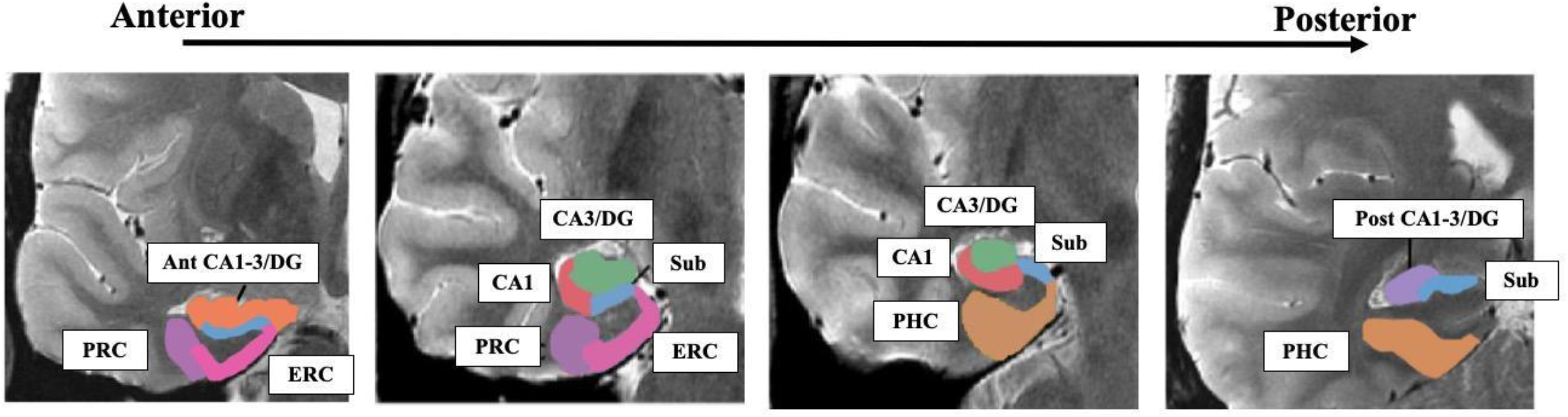
ROI masks were generated by manually tracing on individual anatomical T2-weighted images using the coronal slices that were perpendicular to the main axis of the hippocampus. Masks for hippocampal subfields, including anterior CA1-3/DG (Ant CA1-3/DG), CA3/DG, posterior CA1-3/DG (Post CA1-3/DG), subiculum (Sub), and CA1, were combined to generate whole hippocampal masks.

**Figure S2.**
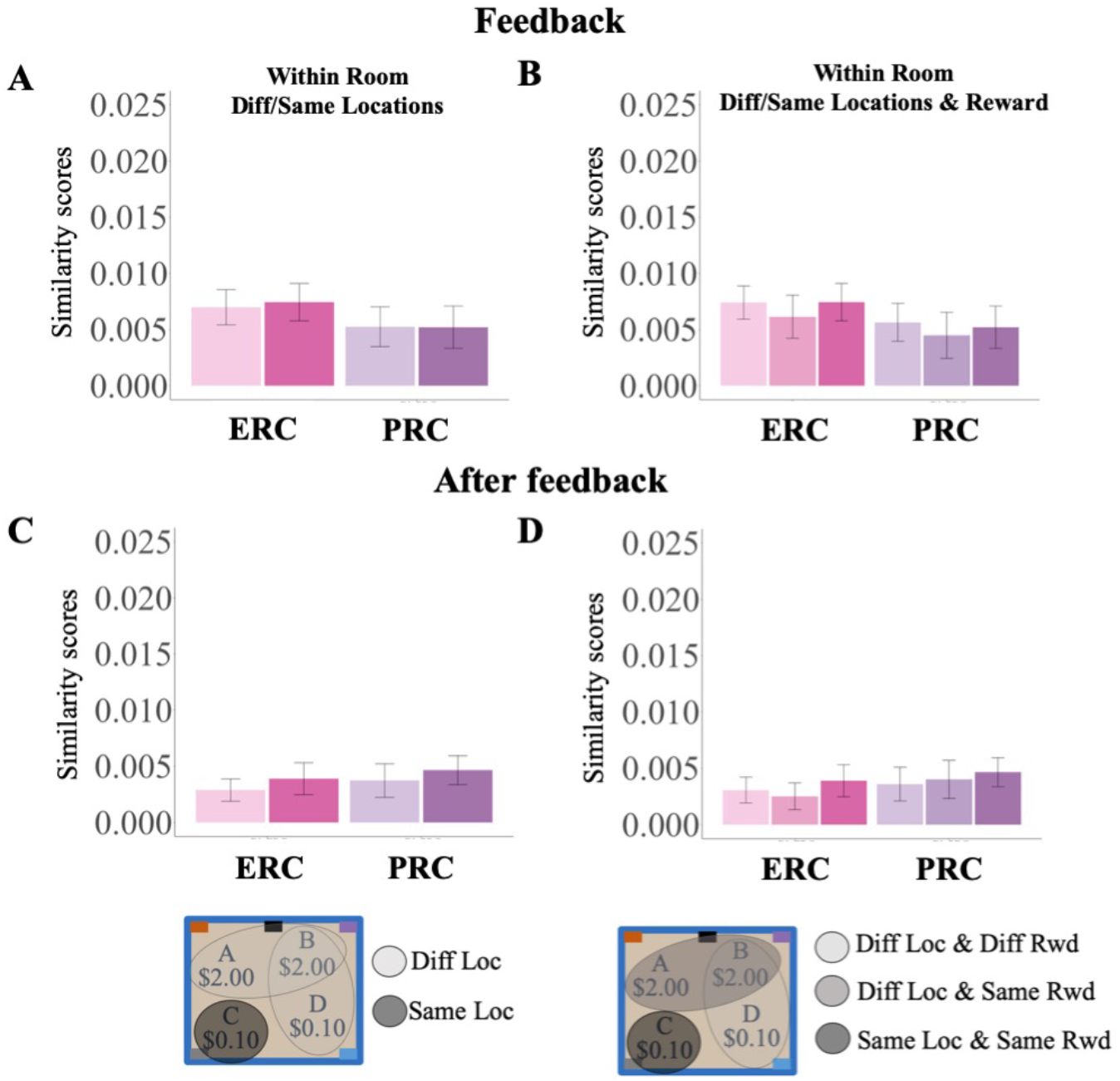
Pattern similarity scores in PRC and ERC for within-room different/same locations and within-room different/same location and reward during the feedback period (A/B) and after-feedback period (C/D). For the plots of within-room different/same locations (A/C), the light color bars represent similarity scores for different locations and the dark color bars represents similarity scores for the same locations. For the plots of within-room different/same locations and reward (B/D), the light color bars represent similarity scores for different locations, the intermediate color bars represent similarity scores for different locations and same reward values, and the dark color bars represent similarity scores for the same location and same reward. Conditions of the bars are demonstrated in the legend: the legends use topographic view of the square room with blue walls as an example similar to Figure 1A. The gray circles in the topographic view of the room are example conditions. Error bars: across-subject standard error of the mean.

**Figure S3.**
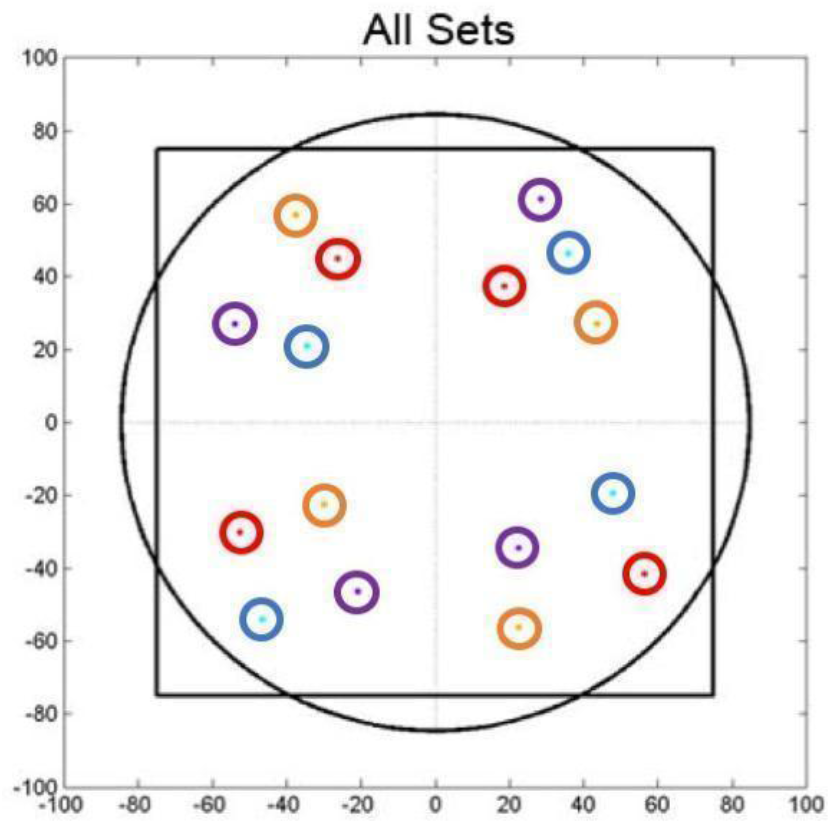
Four candidate sets of the goal locations used in the study: one color represents one set of the locations. Black square and circle lines represent the walls of the square and circular rooms. Dots inside the circles are the centers of the goal locations and the circles represent 5 a.u. radius from the center of the goal location. Movement to within the correct circle was considered as the participant having arriving at the goal location (see Detailed Methods).

### Supplementary text

Below are the full statistical results for global context, reward values, and across room same/different goal quadrants analyses for each task period.

#### Cue period (MTL ROIs × conditions)

same/different room:

condition: F(1,14) = 0.82, p > 0.38; ROIs × conditions: F(3,42) = 0.55, p > 0.64).

*Within room:*

same/different goal locations: condition: F(1,14) = 0.03, p > 0.84; ROIs × conditions: F(3,42) = 1.35, p > 0.27).

same/different reward values: condition: F(1,14) = 2.30, p > 0.15; ROIs × conditions: F(3,42) = 1.22, p > 0.31).

same/different goal locations and same/different reward values: condition: F(1,14) = 2.79, p > 0.07; ROIs × conditions: F(3,42) = 0.90, p > 0.49).

*Across room:*

same/different goal quadrants: condition: F(1,14) = 0.04, p > 0.94; ROIs × conditions: F(3,42) = 0.06, p > 0.97).

same/different reward values: condition: F(1,14) = 0.08, p > 0.77; ROIs × conditions: F(3,42) = 3.02, p > 0.040; post-hoc paired t-test: ps > 0.80; Bonferroni corrected).

#### Held period (MTL ROIs × conditions)

same/different room:

condition: F(1,14) = 0.90, p > 0.35; ROIs × conditions: F(3,42) = 0.58, p>0.63).

*Within room:*

same/different goal locations: condition: F(1,14) = 1.79, p > 0.20; ROIs × conditions: F(3,42) = 0.70, p > 0.55).

same/different reward values: condition: F(1,14) = 1.16, p > 0.29; ROIs × conditions: F(3,42) = 1.35, p > 0.27).

same/different goal locations and same/different reward values: condition: F(2,28) = 0.91, p > 0.41; ROIs × conditions: F(6, 84) = 0.85, p > 0.53).

*Across room:*

same/different goal quadrants: condition: F(1,14) = 0.47, p > 0.50; ROIs × conditions: F(3,42) = 1.90, p > 0.14).

same/different reward values: condition: F(1,14) = 1.11, p > 0.31; ROIs × conditions: F(3,42) = 1.89, p > 0.14).

#### Feedback period (MTL ROIs × conditions)

same/different room:

condition: F(1,14) = 0.10, p > 0.75; ROIs × conditions: F(3,42) = 0.88, p > 0.45).

*Within room:*

We have reported the analyses in the main text.

*Across room:*

same/different goal quadrants: condition: F(1,14) = 0.03, p > 0.87; ROIs × conditions: F(3,42) = 0.47, p > 0.70).

same/different reward values: condition: F(1,14) = 0.01, p > 0.90; ROIs × conditions: F(3,42) = 2.14, p > 0.10).

#### After-feedback period (MTL ROIs × conditions)

conditions: same/different room:

condition: F(1,14) = 0.19, p > 0.66; ROIs × conditions: F(3,42) = 0.69, p>0.56).

*Within room:*

same/different reward values: condition: F(1,14) = 0.50, p > 0.48; ROIs × conditions: F(3,42) = 2.76, p > 0.05).

same/different goal locations and same/different reward values: condition: F(2,28) = 2.13, p > 0.13; ROIs × conditions: F(6,84) = 1.72, p > 0.12).

*Across room:*

same/different goal quadrants: condition: F(1,14) = 0.029, p > 0.86; ROIs × conditions: F(3,42) = 1.23, p > 0.31).

same/different reward values: condition: F(1,14) = 0.01, p > 0.91; ROIs × conditions: F(3,42) = 1.99, p > 0.13).

